# A benzene-mapping approach for uncovering cryptic pockets in membrane-bound proteins

**DOI:** 10.1101/2020.04.04.025163

**Authors:** Lorena Zuzic, Jan K Marzinek, Jim Warwicker, Peter J Bond

## Abstract

Molecular dynamics (MD) simulations in combination with small organic probes present in the solvent have previously been used as a method to reveal cryptic pockets that may not have been identified in experimental structures. We report such a method implemented within the CHARMM forcefield to effectively explore cryptic pockets on the surfaces of membrane-embedded proteins using benzene as a probe molecule. This relies on modified non-bonded parameters in addition to repulsive potentials between membrane lipids and benzene molecules. The method was tested on part of the outer shell of the dengue virus (DENV), for which research into a safe and effective neutralizing antibody or drug molecule is still ongoing. In particular, the envelope (E) protein, associated with the membrane (M) protein, is a lipid membrane-embedded complex which forms a dimer in the mature viral envelope. Solvent mapping was performed for the full, membrane-embedded EM protein complex and compared with similar calculations performed for the isolated, soluble E protein ectodomain dimer in solvent. Ectodomain-only simulations with benzene exhibited unfolding effects not observed in the more physiologically relevant membrane-associated systems. A cryptic pocket which has been experimentally shown to bind *n*-octyl-β-D-glucoside detergent was consistently revealed in all benzene-containing simulations. The addition of benzene also enhanced the flexibility and hydrophobic exposure of cryptic pockets at a key, functional interface in the E protein, and revealed a novel, potentially druggable pocket that may be targeted to prevent conformational changes associated with viral entry into the cell.

**GRAPHICAL ABSTRACT:** 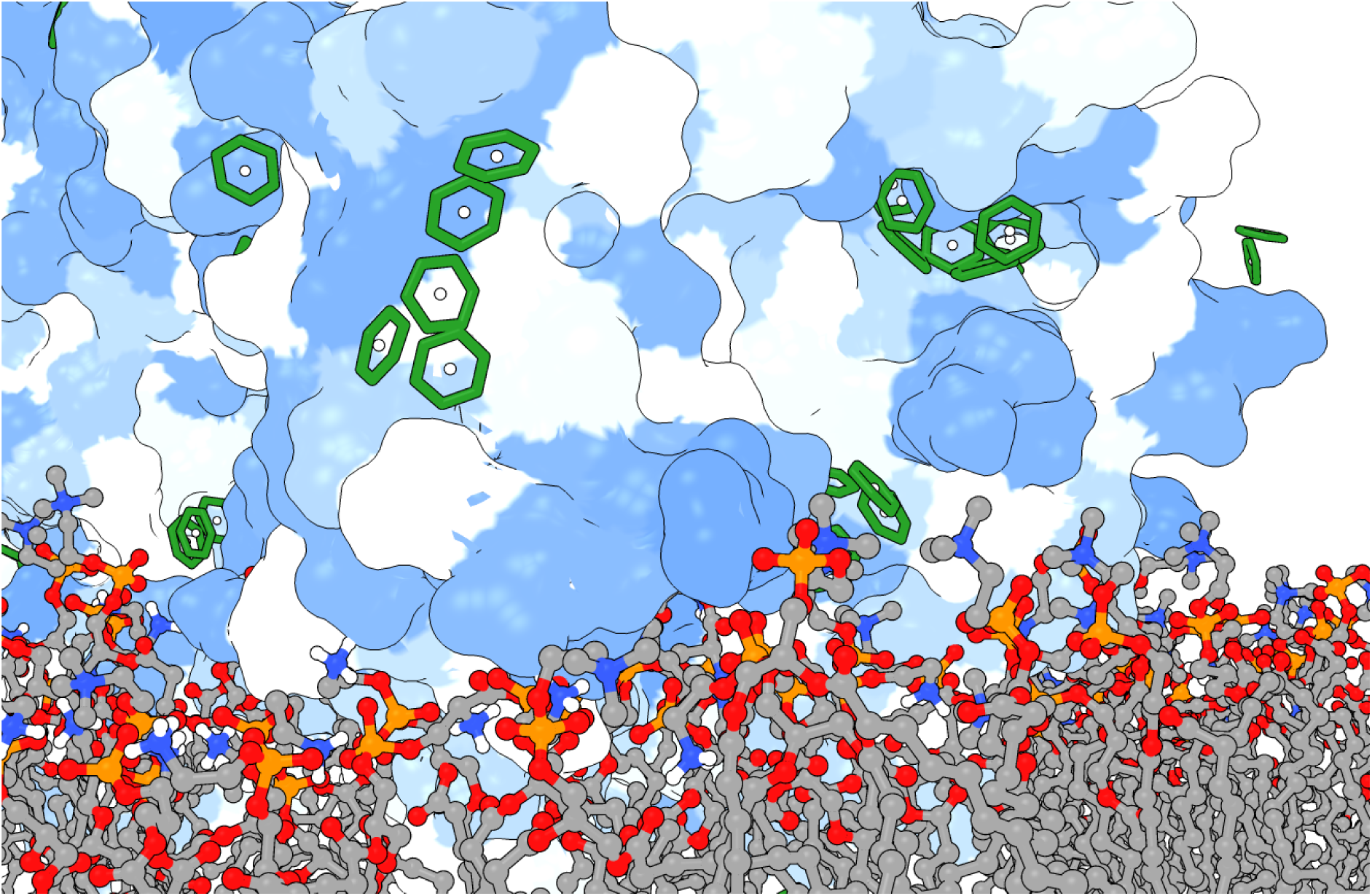

## 1 INTRODUCTION

Drug development has long focussed on proteins as the principal druggable targets, mainly due to their potential to exhibit functional modulation as a response to specific binding of a drug molecule^1,2^. The development of protein-specific drugs hinges upon the presence of functional drug-binding hotspots on the protein surface or within the protein interior^3^. Hotspots are regarded as protein regions whose physical appearance (often pocket-like) and chemical properties inordinately contribute to a favourable binding-free energy of a ligand^4^. As such, hotspot detection and characterisation is a desirable prerequisite for rational drug design. It has been shown experimentally that small organic molecules do not indiscriminately cover the protein surface, but instead have a tendency to bind to hotspots^5^. This facet has been used to computationally detect hotspots using simple organic molecules (fragments or probes) on both rigid and dynamic proteins of interest^6^. The majority of probe-based research has so far been focussed on hotspot detection on static protein structures using grid- or minimisation-based methods^7–9^. These methods consider proteins as rigid bodies with well-defined drug-binding sites present in the crystal structure. The reality, however, is that a large number of proteins of interest do not typically exhibit any obvious hotspots suitable for specific drug-binding, according to their static experimental structures. This, in turn, makes the pursuit of targeted druggability based solely on crystal or NMR structures inadequate for a non-negligible portion of researched proteins^10,11^.

MD simulation is a computational method that calculates the likely movements of a macromolecule and its immediate environment by applying Newton’s equations of motion and force field-defined parameters to all atoms present in the system. In recent years, high performance computing has enabled long-timescale, biologically meaningful sampling of large macromolecular systems^12^, while continually improved and refined force fields provide better accuracy of the simulated data^13^. This, in turn, has made possible the simulation of “rare events” and the exploration of less-populated protein conformational states for potential drug-binding hotspots^14,15^. However, conventional MD simulations, even when paired with long simulation timescales or improved sampling methods, are not yet routinely able to reliably detect the full repertoire of transient binding pockets on protein targets^16^. Cryptic pockets are usually hydrophobic in nature, making their opening and exposure to the highly polar water environment energetically unfavourable, and thus under-sampled by the simulated system^17,18^. Typically, cryptic pockets are reliably revealed only in the presence of a ligand which binds inside the pocket and locks it in an open conformation long enough for it to be detected^19,20^. Simulations using small organic probes, either as homogenous or mixture co-solvents^18,21–23^, have proven useful for the detection and characterisation of cryptic pockets on a whole range of therapeutically relevant proteins^14,18,24^. Solvent-mapping MD methods employ a range of probe concentrations. High probe concentrations allow for more efficient mapping of cryptic pockets and other surface ligand-binding sites, but they can also result in aggregation and phase-separation of hydrophobic probes from the bulk solvent. The issue has been addressed by introducing artificial repulsion terms between the individual probe molecules^21^ or by lowering the overall probe concentration of the system^18^.

Addition of small probe molecules to the system also presents a major issue if a protein of interest is membrane-bound, as the probes have a propensity to migrate into the hydrophobic lipid environment and remain there during the course of the simulation^25^. Therefore, even though large portions of potential drug targets are membrane proteins^26^, probe-based MD simulations have seldom been applied to systems containing a lipid bilayer. Probe sequestration in protein-membrane systems was successfully inhibited by Gorfe and co-workers, by adding non-bonded repulsive interaction potentials between the polar probes and lipid tail methylene group carbon atoms, as demonstrated for K-Ras^23,27^. In this method, the placement of the lipid repulsive points below the glycerol backbone allowed the probes to interact with the headgroup portion of the phospholipid bilayer. A relatively small (0.7 - 1.0 nm) repulsion distance successfully excluded the probes containing at least one tetrahedral moiety from the hydrophobic portion of the membrane. For the most part, the probes used in this method were soluble in water and exhibited no solute partitioning; isobutane was prevented from self-aggregating by introducing short range LJ repulsions on a centrally placed carbon atom^21^. Nevertheless, the applicability of the method and the proposed parameters to a benzene molecule remains to be addressed, particularly with regards to aggregation, depth of entry into the membrane, and the stringency of the repulsion distance parameter.

Using pocket mapping for viral proteins is an attractive area of study with the potential to utilize transient pocket discovery for targeted antiviral drug design and for identifying conserved but hidden antibody-binding epitopes^28–30^. Flaviviruses are a family of enveloped, single-stranded RNA viruses that include a range of vector-borne human pathogens such as DENV, Zika virus, yellow fever virus, or Japanese and tick-borne encephalitis virus^31^. These viruses present a major health burden on a global scale, especially since climate change is expanding the range of virus-carrying mosquitoes previously contained predominantly to the tropical region^32,33^. DENV, in particular, is responsible for 390 million cases worldwide, 96 million of which are symptomatic^34^. The DENV particle consists of a nucleocapsid encapsulated in a spherical envelope; the surface is comprised out of 90 orderly packed, icosahedral E protein dimers anchored to a viral membrane. Underneath each dimer ectodomain, there are two smaller M proteins which are hidden from the surface in the mature viral particle^35^. In the extracellular environment, the virus mostly occurs in a mature conformation where the only exposed structural element is the E protein ectodomain, making it an ideal binding target for specific drugs or antibodies that have the capacity to inhibit the virus in the early stages of its life cycle^30,36^.

DENV exhibits significant structural heterogeneity, caused not only by subtle sequence variability or maturation deformities, but also by environmental factors such as temperature and pH changes, typically encountered during passage through host micro-environments during the viral life cycle^37–40^. This adds to the complexity of developing vaccines whose antibodies can effectively neutralize all viral strains without inadvertently causing antibody-dependent enhancement (ADE), which can lead to the worst forms of dengue associated disease^41,42^. Therefore, finding a common and biologically relevant binding site on the E protein and targeting it with a small drug-like molecule might be the preferred treatment approach that sidesteps the issues associated with ADE whilst still being capable of acting in an inhibitory manner, potentially across a range of flaviviral strains.

In this study, we present an MD-based solvent-mapping method for cryptic pocket discovery which is applicable to systems containing lipid membranes. Building on the use of a benzene probe strategy^21^ to prevent aggregation, an additional set of repulsive interactions between the probes and key lipid sites are defined to disfavour sequestration of benzene within the membrane interior. Benzene molecules are therefore more likely to map the protein surface, in spite of the presence of the lipid bilayer. We systematically approach the optimization of a membrane-probe repulsive term by testing it for a range of non-bonded Lennard-Jones parameters and its placement on several heavy atoms along the lipid molecule. Finally, we evaluate this approach by exploring cryptic pockets on the E protein dimer of dengue serotype 2 (DENV-2) embedded within a biologically relevant lipid membrane environment, and show that the method is efficient in mapping existing pockets, but also reveals a novel pocket in a functionally significant region of the protein complex.

## 2 METHODS

### 2.1 BENZENE PARAMETERS

Benzene molecule partial charges were taken from the initial phenylalanine aromatic ring parameters specified in the CHARMM36 force field^43^ and adjusted for the uniform charge distribution across all carbon atoms. Exclusions for all intermolecular benzene-benzene interactions were applied so as to allow for denser packing of the probes on the protein surface. In order to avoid clustering of benzene probes, an uncharged dummy atom was centrally placed inside each benzene ring and constrained at a 0.1375 nm distance from each of the heavy atoms using the LINCS algorithm^44^. To prevent probe aggregation, a non-bonded repulsive term was defined between dummy atoms via the Lennard-Jones (LJ) potential using a small value for the maximum depth of the potential well, ε = 0.008 kJ mol-1, and the distance at which inter-particle potential reaches zero, σ = 0.45 nm. These benzene parameters have been optimized for cryptic pocket probe-mapping in soluble proteins, as described in detail in Soni^*45*^.

### 2.2 ASSESSMENT OF MEMBRANE PROPERTIES

Initially, three membrane systems with different lipid compositions were simulated in the absence of benzene in order to establish mean membrane properties including lateral lipid headgroup distance, and hence, possible points of entry of the probes into the bilayer. We chose two zwitterionic homogenous membranes, 1-palmitoyl-2-oleoyl-phosphatidylcholine (POPC) and 1-palmitoyl-2-oleoyl-phosphatidylethanolamine (POPE) bilayer, and one heterogeneous membrane containing a combination of POPC, POPE, and anionic 1-palmitoyl-2-oleoyl-phosphatidylserine (POPS), in a 6:3:1 ratio (Fig. 1). The composition of the heterogeneous membrane was based on lipidomics data available for the DENV-2 virus grown in a C6/36 mosquito cell line^46^.

**Figure 1.**
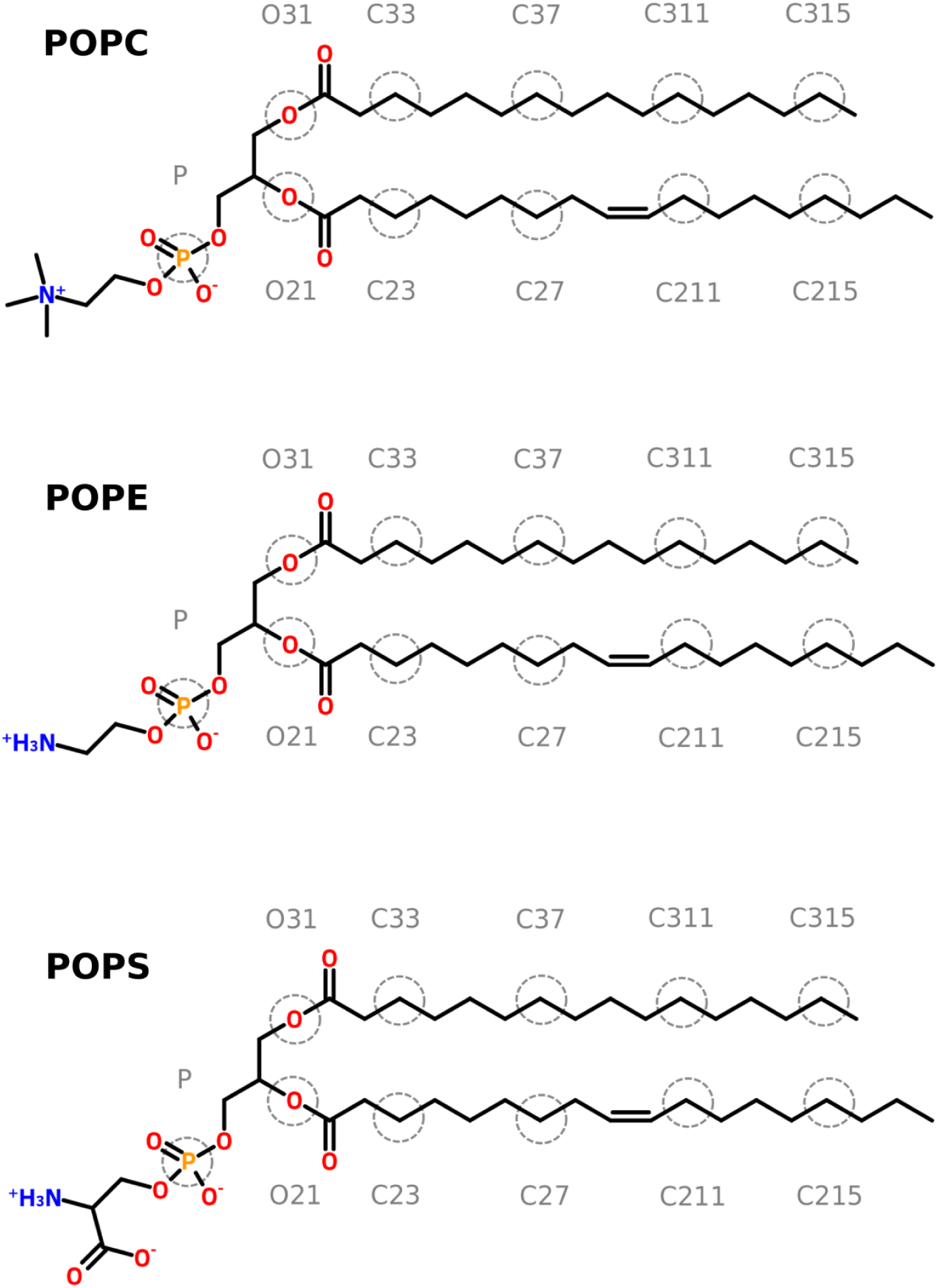
Points of repulsion on the phospholipids present in the viral membrane. We chose the atoms that were present in most phospholipids, evenly distributed down the lipid tail, and present in both oleolyl and palmitoyl fatty acid chains. Because of their heterogeneity between lipids, headgroup atoms were avoided as points of repulsion.

### 2.3 REPULSION TERMS AND FORCE FIELD MODIFICATION

Repulsive terms designed to prevent the probe molecules from entering the lipid bilayer were defined as non-bonded interactions between the benzene dummy atoms and specific atoms of each membrane lipid. The LJ term ε was set to a near-zero value of 0.008 kJ mol^-1^, and the σ_B-L_ – value (subscript referring to the benzene-lipid interactions) was newly parametrized in an iterative fashion to respond correctly to benzene molecules entering the membrane through lateral gaps between the lipids.

The σ_B-L_ parametrization step was performed on systems containing a heterogeneous DENV membrane and 50 benzene molecules placed in bulk solvent (an arbitrarily chosen high concentration, corresponding to a ∼1.9 M solution). A range of σ_B-L_ values (0.8 – 1.4 nm) was selected based on lateral lipid areas and the corresponding distances observed between neighbouring lipids in the probe-free membrane systems. The repulsive potential was initially introduced between each lipid phosphorus atom and each benzene dummy atom. The goal was to establish the repulsive interactions with a threshold σ_B-L_ value that is able to keep the probes outside the bilayer, whilst reducing possible interference with probe-binding to a lateral portion of a protein close to the membrane.

After establishing the optimal σ_B-L_ term, the effectiveness of the benzene repulsion was tested against six different combinations of repulsion points on the lipid molecule, chosen in a way so that they are positioned at different depths below the membrane surface (P only, or atom pairs including O21/O31, C23/C33, C27/C37, C211/C311, and C215/C315; atom names are shown in Fig. 1). We used the same heterogeneous DENV membrane system with 50 benzene probes in the system and varied only the placement of the repulsion point down the lipid tails.

### 2.4 SYSTEM PREPARATION

#### 2.4.1 Pure membrane simulations

All membranes were created using CHARMM-GUI software^47^. Each membrane contained around 80 lipids and were approximately 5 x 5 nm^2^ in size. We used Gromacs 2018 package^48^ to solvate the simulation box with ∼3,000 TIP3P water molecules^49,50^ and to add NaCl salt in a neutralizing 0.15 M concentration. In simulations with benzenes, 50 probe molecules were randomly placed outside of the bilayer in bulk solvent.

#### 2.4.2 DENV-2 envelope – membrane (EM) protein dimer

Verification of the benzene-mapping method was performed on a mature DENV-2 EM protein dimer^35^ (PDB: 3J27) which was inserted in a dengue-specific mixed POPC / POPE / POPS membrane. Based on the results from the membrane-only simulations, we used force fields with two different points of repulsion (P and O21/O31 atoms) and tested them separately (Fig. 1; Table 1). Initial protein alignment in the *xy*-plane was performed using the PPM Server^51^, followed by a membrane building step in CHARMM-GUI^52^. With the exception of the benzene-free control, we added 332 benzene molecules (corresponding to a 0.4 M probe solution) at randomly chosen initial positions and at a minimum distance of 0.5 nm from both protein and the membrane. The systems were then solvated in ∼30,000 TIP3P water molecules and 0.15 M NaCl, neutralizing the overall system charge.

**Table 1.**
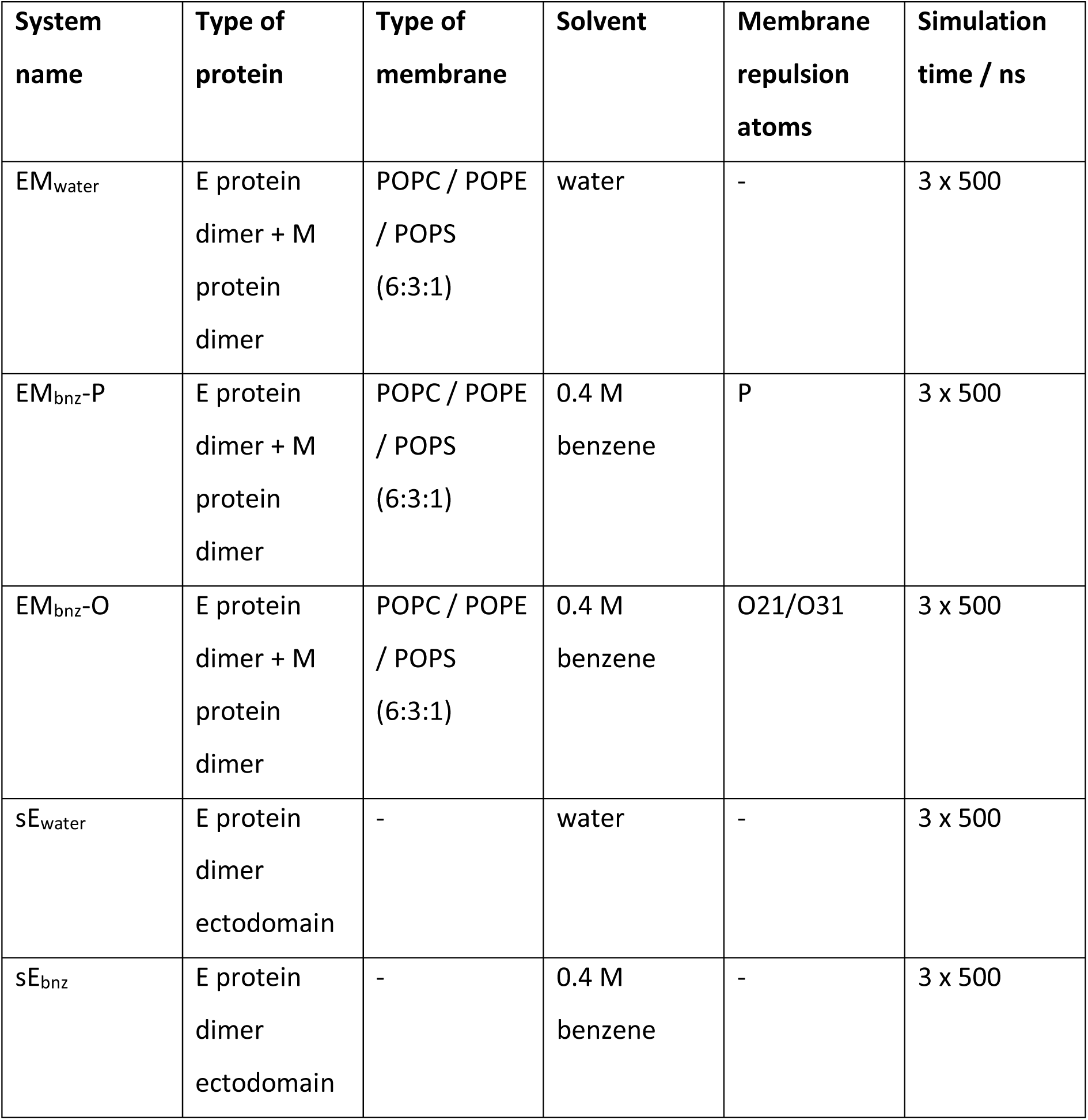
Simulation systems containing DENV-2 proteins against which the method was verified. The effectiveness of cryptic pocket mapping on a membrane-bound EM dimer was compared with the existing method for a membrane-free system (in this case, the sE ectodomain dimer). Both P and O21/O31 lipid atoms were used as points of repulsion in separate simulations in the presence of membrane. Probe-free controls were run for both EM and sE system types.

| System name | Type of protein | Type of membrane | Solvent | Membrane repulsion atoms | Simulation time / ns |
| --- | --- | --- | --- | --- | --- |
| EM<sub>water</sub> | E protein dimer + M protein dimer | POPC / POPE / POPS (6:3:1) | water | - | 3 x 500 |
| EM<sub>bnz</sub>-P | E protein dimer + M protein dimer | POPC / POPE / POPS (6:3:1) | 0.4 M benzene | P | 3 x 500 |
| EM<sub>bnz</sub>-O | E protein dimer + M protein dimer | POPC / POPE / POPS (6:3:1) | 0.4 M benzene | O21/O31 | 3 x 500 |
| sE<sub>water</sub> | E protein dimer ectodomain | - | water | - | 3 x 500 |
| sE<sub>bnz</sub> | E protein dimer ectodomain | - | 0.4 M benzene | - | 3 x 500 |

#### 2.4.3 DENV-2 envelope protein ectodomain (sE)

We tested the efficacy of our method by comparing it to a system containing a DENV-2 E protein ectodomain dimer (residues 1 to 395) missing the transmembrane domains and stem regions (residues 396 to 495), with no membrane present. The system was placed in a truncated dodecahedron box and solvated in 0.4 M benzene^45^, 0.15 M NaCl, and ∼120,000 TIP3P water molecules. We also ran a probe-free control on the ectodomain dimer in water.

### 2.5 SIMULATION PROTOCOLS

All simulations were performed using modified CHARMM36 force field^43^ and Gromacs 2018 simulation software^48^. Systems were minimized using the steepest descent algorithm. A temperature of 310 K and pressure of 1 atm was maintained using the velocity-rescaling thermostat^53^ and Parrinello-Rahman barostat^54^, respectively. A 2 fs integration time step was paired with LINCS constraints on all bonds associated with hydrogen atoms along with those constraining the dummy atom at the centre of each benzene probe molecule^44^. The Particle mesh Ewald (PME) method was used for describing long-range electrostatic interactions^55^. The short-range van der Waals cutoff and real-space cutoff for PME were both set to 1.2 nm, with included potential-shift Coulomb and force-switch van der Waals modifiers with a 1.0 nm cut-off distance.

All position restraints applied to the system components in equilibrium simulations were defined with a force constant of 1,000 kJ mol^-1^ nm^-2^ in all three dimensions. For membrane-only systems, an initial *NVT* equilibration was run for 50 ps with position restraints on all heavy membrane and probe atoms. In order to allow for sufficient membrane relaxation, a longer subsequent equilibration of 100 ns was run with position restraints applied to benzene molecules only, followed by an unrestrained 100 ns production run.

EM dimer systems embedded within a membrane were equilibrated for 1 ns in the *NVT* ensemble with position restraints applied to all heavy atoms of protein, membrane, and benzene components. Subsequently, *NPT* equilibration was run for 100 ns with position restraints applied only to benzene carbon atoms along with the E and M protein stem and transmembrane helices. These stem and transmembrane helix restraints were essential for retaining a membrane curvature observed in the respective cryo-EM structure^35^, and were maintained during subsequent 500 ns-long production runs. Similarly, sE (ectodomain-only) systems involved an initial 1 ns *NVT* run with position-restrained protein and benzene components, followed by a 100 ns *NPT* equilibration with position restraints applied only to benzenes, and finally, an unrestrained 500 ns production run. All simulations are listed in Table 1. All production simulations were run in three independent repeats, resulting in 1.5 μs of total simulation sampling for each system.

### 2.6 SIMULATION ANALYSIS

Key protein properties, including density, solvent-accessible surface area^56^, or secondary structure content^57^, were calculated in the Gromacs analysis software suite^48^. Membrane properties were analysed using FatSlim tools^58^. In order to determine lateral gaps between the lipids, phosphorus atom coordinates of each lipid in an *xy* plane were mapped onto a Voronoi surface such that each Voronoi area (representing the occupancy of a single lipid in a leaflet) had a determined, finite number of lipid neighbours (Fig. S1). The maximum distance between a single lipid and its neighbours was then calculated for all (non-periodic boundary bordering) lipids and plotted as an average across a simulation trajectory.

Detection of protein pockets was carried out using the MDPocket software^59^, which allows for cavity mapping across the dynamic protein trajectory and accounts for the stability of a pocket by associating it with a frequency of its detection during the course of the simulation. We considered only the pockets mapped onto the ectodomain portion of the protein. In addition, we excluded pockets that were consistently present across all simulations for the majority of each trajectory, as they were considered unchanging structural features of the protein, rather than cryptic pockets. Instead, we focussed on the regions of the protein which manifested the biggest change from the water-only simulations, and which had sufficient hydrophobic character capable of binding the benzene probes. The efficacy of the designed probe-mapping method was established by comparing known and novel pockets found in the EM simulations with the membrane-free sE systems. Protein electrostatics maps were calculated using the APBS software package^60^.

## 3 RESULTS AND DISCUSSION

### 3.1 PROTEIN-FREE MEMBRANE SIMULATIONS

The flat, planar shape of the benzene ring allows it to enter the membrane through the lateral gaps between headgroups, even after adding short-range repulsive forces between the probe and the membrane (previously described by Prakash *et al*.^*27*^). The authors used a LJ σ_B-L_ value of 0.7 nm for their simulations, which is adequate for bulkier or branched probe molecules such as isopropanol, but is insufficient for benzene which rapidly partitions into the hydrophobic layer of the membrane. In order to assess the size of the lateral gaps between lipid headgroups, as well as their variability depending upon the membrane lipid composition, we ran three probe-free 100 ns membrane simulations with differing lipid contents (pure POPC, pure POPE, or a heterogeneous mixture of POPC / POPE / POPS in a 6:3:1 ratio). In all three cases, the maximum gap between any lipid and its nearest neighbours varied between a minimum of 1.1 nm and a maximum of 1.25 nm (Fig. S2), suggesting that the σ_B-L_ term, which determines the range of non-bonded interactions, had to be increased in order to prevent benzene from entering the membrane through the gaps. We subsequently ran eight 100 ns simulations of the heterogeneous membrane with benzene and with σ_B-L_ values in a range that encompassed the previously calculated maximum distance values (Fig. 2A). The results show that the probe is able to enter the membrane if σ_B-L_ is below a value of 1.2 nm. If greater than or equal to 1.2 nm, benzene molecules remain, for the most part, in the aqueous environment of the simulated system.

**Figure 2.**
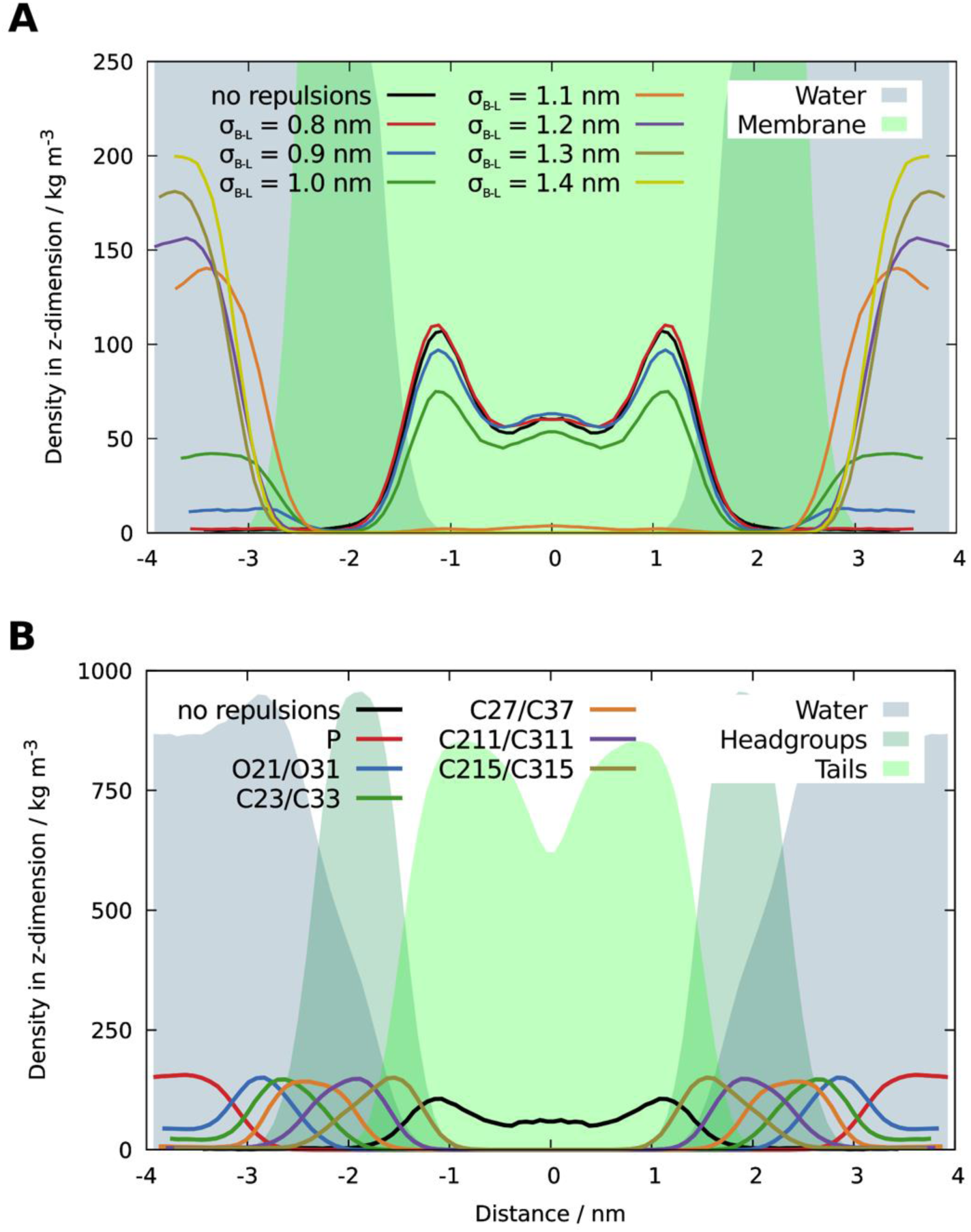
Density distribution of system components including benzene probes in the *z*-dimension. The density depends on (A) the σ_B-L_ value defined for the non-bonded repulsive term between the membrane and benzene probes, (B) the placement of repulsive points on the lipid tails. Solid lines show symmetrized distribution of benzene probes. Transparent surfaces indicate membrane and water components of the system.

Specifying the repulsive potentials such that they operate over a longer range than is necessarily required might improve their effectiveness in keeping the probes outside the membrane core. However, this may also introduce the unintended effect of preventing the probes from binding to the potentially druggable portions of the protein which are close to the membrane, but not embedded within it. We therefore opted for the σ_B-L_ value of 1.2 nm, since this is the shortest repulsive distance that is able to adequately prevent benzene from sequestering into the bilayer.

In addition to testing different σ_B-L_ values, we also considered a selection of membrane lipid atoms which could be defined as points of repulsion against probe molecules. We avoided using headgroup atoms because of their heterogeneity, dependent upon the lipid type, which would make generalizing the approach more difficult. As shown in Figure 2B, repulsion points positioned lower down the lipid chain also permit benzene to occupy the bilayer at increasingly greater depths. Depending on the force field modification, benzene can predominantly be distributed in water (atoms P, O21/O31), the hydrophilic headgroup portion (atoms C231/C33, C27/C37, C211/C311) or the hydrophobic tail portion (atoms C215/C315) of the membrane. A large number of hydrophobic probes interacting with lipids can have an effect upon membrane properties compared to the innate, benzene-free state (Fig. S3). Retaining key equilibrium membrane characteristics close to those observed in a probe-free environment may be achieved by reducing the number of contacts between lipids and benzene molecules, regardless of if the contacts are with the hydrophilic or the hydrophobic portion of the membrane. Additionally, by reducing the ratio of lipid-associated benzene molecules to a minimum, this ensures that most probe molecules are free to interact with the protein, making the benzene mapping process and exposure of cryptic pockets overall more effective.

### 3.2 DENV-2 E PROTEIN DIMER SIMULATIONS

#### 3.2.1 Solvent-accessibility of the E protein dimer ectodomain

Simulations involving a protein component were performed on systems with membrane-embedded E and M protein dimers (EM), as well as ectodomain dimers without the presence of a membrane (sE). The EM complex was set up in probe-free conditions (EM_water_) or with benzene and lipid repulsion points placed on P (EM_bnz_-P) or O21/O31 lipid atoms (EM_bnz_-O; snapshot shown in Fig. 3A). The sE complex was simulated without benzene probes (sE_water_), or with the addition of benzene (sE_bnz_; snapshot shown in Fig. 3B).

**Figure 3.**
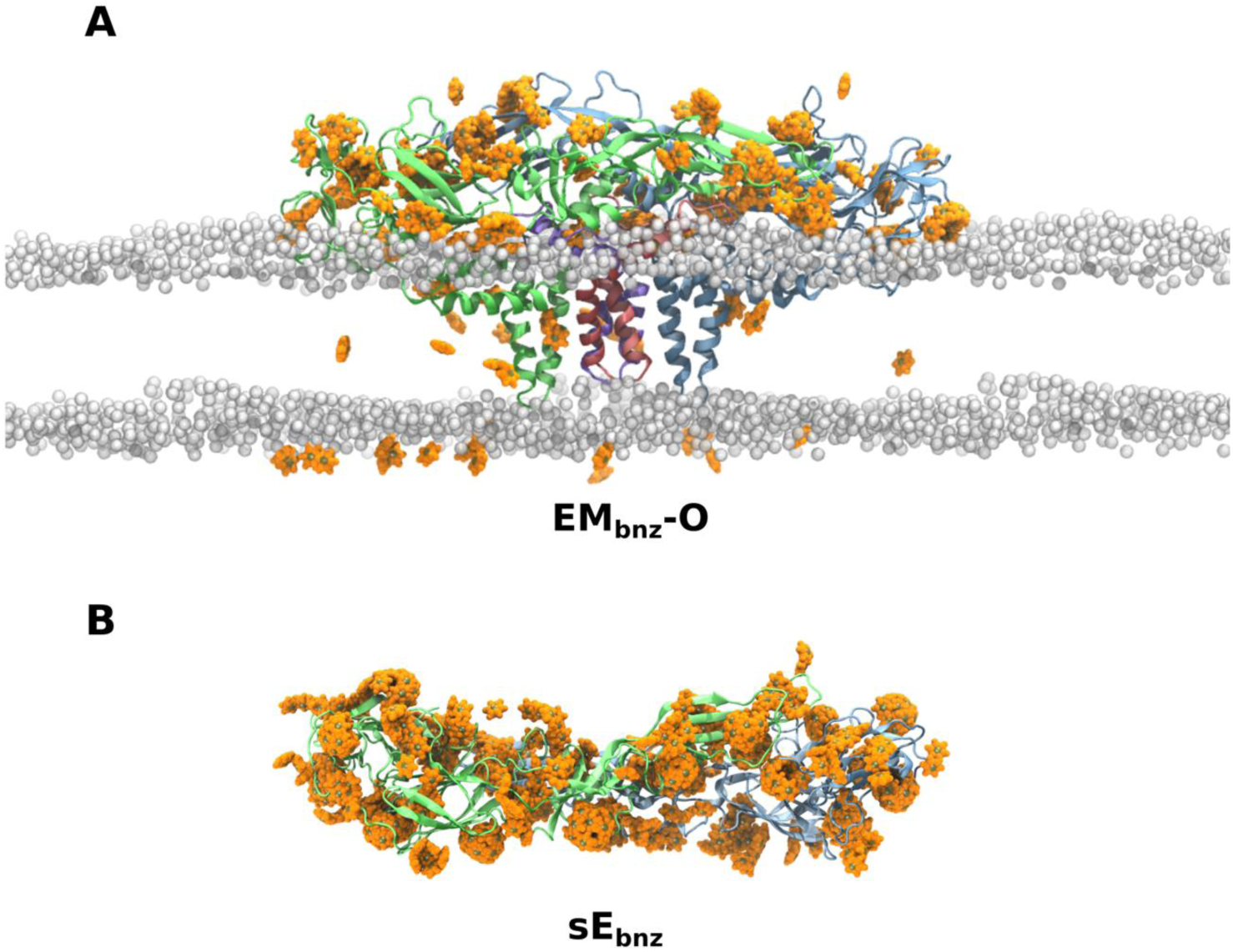
Side view of the representative snapshots for two simulated systems. (A) is the EM_bnz_- O system which includes the EM dimer and benzene probes in the presence of a membrane, and (B) is the sE_bnz_ system, which contains the ectodomain-only E protein dimer and benzene without the membrane present. For clarity, only membrane phosphorus atoms and benzene probes close to the protein or the membrane are shown. Protein is shown in cartoon representation, E chains coloured in green and blue and M chains in pink and purple. Benzenes are shown as orange van der Waals spheres with dummy atoms indicated in tan.

The success of probe-mapping methods is customarily measured by its ability to increase solvent-accessible surface area (SASA) of the protein, which is partly a result of the favourable exposure of cryptic pockets to the interacting solvent^61^. The hydrophobic character of a probe increases the probability that, upon binding, the pocket – often hidden inside the protein interior – opens up. Simulation systems containing only the E protein ectodomains are likely over-simplified, neglecting the effects of the lipid bilayer as a hydrophobic barrier and as an interaction plane for the lower surface of the protein (Fig 4A). Furthermore, the absence of key system components such as E protein stem regions and transmembrane domains, neighbouring M proteins, along with the lipid membrane allow the solvent to interact with both upper and lower surfaces of the protein, which is in contradiction with the conditions present in the mature viral particle where solvent interacts primarily with the upper surface of the envelope, leaving the lower surface largely unaffected^62^. SASA between the two simulation setups, therefore, is only commensurable for the upper surface portion of the E proteins.

**Figure 4.**
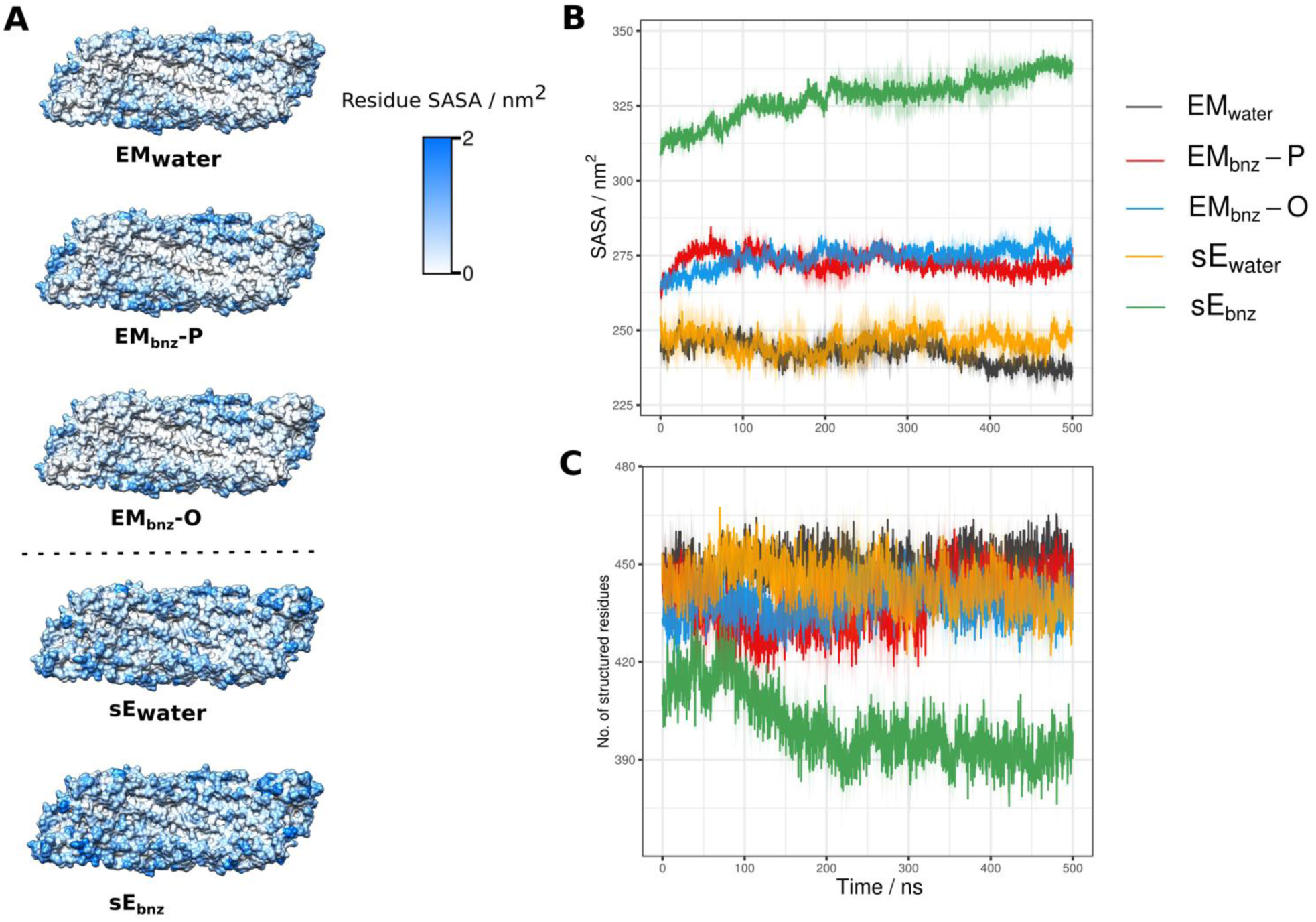
The effect of benzene probes on solvent accessibility and secondary structure evolution. (A) The figure shows surface representations of the residue SASA measured for the underside of each ectodomain dimer. In the virion, this surface is oriented towards the lipid membrane and interior of the virus. SASA of the ectodomain dimer underside is affected by the presence or absence of a membrane. The lower surface of the ectodomain for systems in the presence of a lipid membrane (EM systems) is for the most part inaccessible to the solvent, reflecting the organisational features of the viral envelope. Membrane-free, ectodomain-only simulations (sE systems), on the other hand, have the lower surface fully exposed to the solvent, which is manifested in an overall greater SASA. (B) Evolution of SASA over the course of each simulation measured for the ectodomain upper surface. The sE_bnz_ system had drastically higher SASA which continued to increase during course of the simulation without plateauing. The starting values differ between the systems due to equilibration. (C) Ectodomain structural stability reflected in the number of residues forming secondary structure elements remained consistent across all simulation systems except for sE_bnz_, suggesting that it underwent partial unfolding due to benzene present in the solvent.

Upon first inspect, based on solvent accessibility alone, simulating the system containing the E protein ectodomains in the presence of benzene (sE_bnz_) seems like a superior probe-mapping approach (Fig. 4B). Higher exposure to the solvent indicates that sE_bnz_ system is most likely to present a majority of previously hidden cryptic pockets at the protein surface. The increase of SASA, however, is not entirely a consequence of cryptic pocket exposure. Organic solvent also has a potential disruptive effect on the protein structural stability and may lead to denaturation if the organic component concentration is substantial enough to lead to disorder of the hydrophobic core of the folded protein^63,64^. Figure 4C shows a stark decrease in protein secondary structure for sE_bnz_ when compared to the benzene-free simulation, indicating that the addition of benzene is driving the protein to partially unfold in solution. Denatured states, either partial or complete, are not desirable conformations to be sampled in a simulation as they are unlikely to be present in a functional viral particle. Similarly, hydrophobic pockets detected in a protein in its denatured state, as in sE_bnz_, are less likely to exist in the folded protein and therefore irrelevant as potential drug-binding pockets. In contrast, simulations containing the full protein dimer within the membrane environment (EM_bnz_-P and EM_bnz_-O) retain their structural integrity even when exposed to the same concentration of benzene (Fig. 4C). The increase in SASA coupled with the conservation of secondary structure for these systems suggests that including the complete protein and its complex with the M proteins within the membrane environment is better suited for detection of biologically relevant cryptic pockets, compared to a setup containing only an isolated, soluble portion of the protein.

#### 3.2.2 Benzene interactions with protein and membrane

Benzene used as a probe in our systems has been modified to maximize the effectiveness of cryptic pocket exposure, as previously validated^45^. All interactions between benzene molecules were removed from the force field via introduction of exclusions, meaning that benzene ring atoms are effectively “blind” to the existence of other probe molecules in the system. Additionally, a repulsive dummy atom at the centre of each benzene prevents other benzene dummy atoms from approaching at an interaction distance of less than 0.45 nm. These modifications allow for close packing of probe molecules in a hydrophobic pocket space, whilst at the same time preventing aggregation, or excessive overlap within bulk solvent (Fig. 4).

The majority of benzene probes established transient interactions with the protein ectodomain and remained in its vicinity for the entire duration of the simulations (Fig. 5A). A small fraction of the probe molecules managed to enter the membrane despite the repulsive forces with the central dummy atoms. Notably, the EM_bnz_-O system was better at keeping the benzene outside the bilayer, resulting in a greater number of probe molecules being able to map the protein surface.

**Figure 5.**
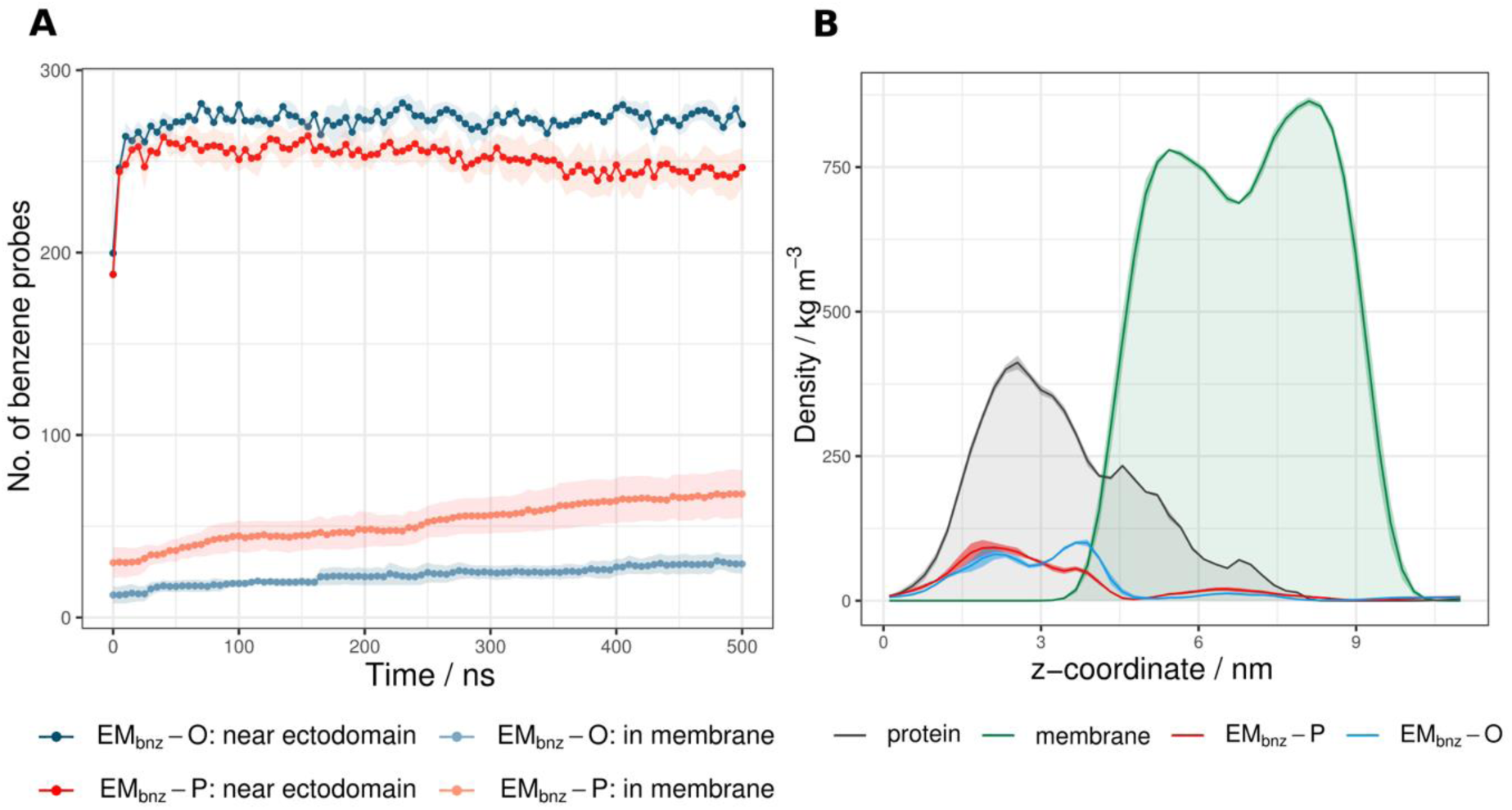
Benzene distribution in the EMbnz simulation systems. (A) Benzene probes detected within a 0.5 nm cutoff distance from the ectodomain and membrane tails. The EM_bnz_-O system overall had more benzene probes in the vicinity of the protein and less probes in the membrane compared to the EM_bnz_-P system. (B) Density distribution of benzene probes with respect to the protein and membrane components. Benzene in the EM_bnz_-P system interacted mostly with the upper E protein ectodomain surface, unlike EM_bnz_-O in which benzene covered both the top and sides of the E protein to a similar extent.

A density profile of system components in the *z*-dimension shows that both EM_bnz_-P and EM_bnz_-O systems have probes that predominantly colocalise with the solvent-exposed “edges” of the protein, featuring two peaks corresponding to the top and to the sides of the ectodomain (Fig. 5B). The EM_bnz_-P system has a repulsion point closer to the membrane surface, resulting in benzene molecules mainly exploring the upper portion of the ectodomain. In contrast, the probe molecules in the EM_bnz_-O system are able to position themselves nearer to the membrane surface, effectively enabling them to map both the upper and side surfaces of the ectodomain. Overall, both systems may be used as a valid setup for benzene mapping of membrane-bound proteins, EM_bnz_-O being moderately more effective in keeping the probes outside the bilayer and exploring all sides of the solvent-exposed portion of the protein.

#### 3.2.3 β-OG pocket detection

Currently, there is only one experimentally confirmed cryptic pocket on the DENV E protein and it is located underneath the *kl* β-hairpin at the domain I - domain II interface^65^ (Fig. 6C). Undetectable in a native DENV-2 crystal structure, it was revealed only upon binding of a single hydrophobic *n*-octyl-β-D-glucoside (β-OG) detergent molecule to its hydrophobic groove, thereby locking the pocket in an open conformation. It is thus desirable that simulation-based methods designed for cryptic pocket discovery should efficiently expose this hydrophobic pocket. Simultaneously, pocket surface area descriptors should closely match the experimental values; otherwise the method risks mapping only a small portion of the pocket or, conversely, a deceptively large pocket due to protein deformation or unfolding. Figure 6A shows that the β- OG pocket was consistently revealed to the solvent in all simulation systems containing benzene. Interestingly, the pocket was also reliably revealed in a benzene-free EM_water_ simulation, suggesting that the presence of a membrane or protein components other than the ectodomain might have long-range effects on the stabilization of the *kl* hairpin in an open conformation. In contrast, the β-OG pocket in the sE_water_ simulation remained firmly closed and hidden from the aqueous environment.

**Figure 6.**
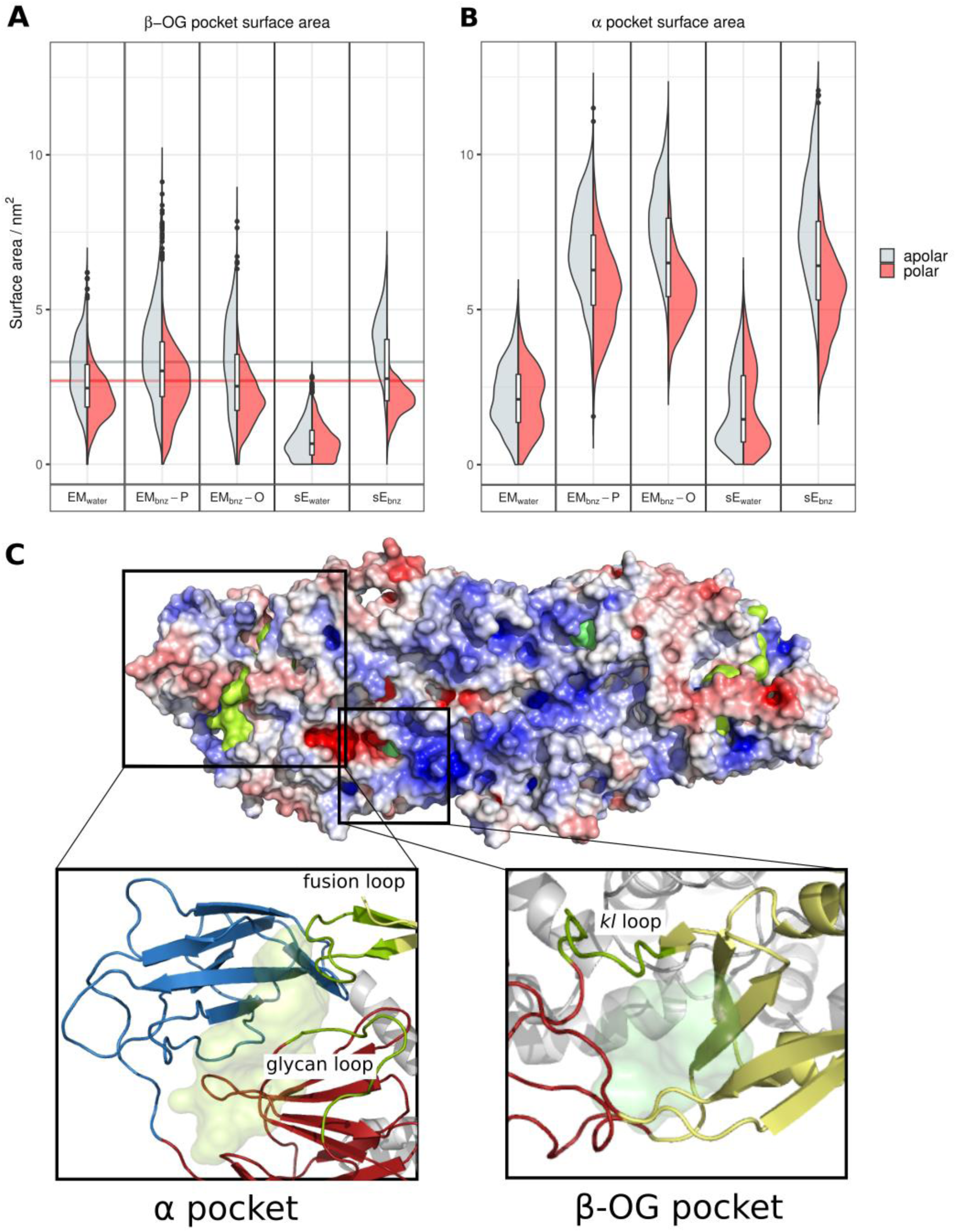
Existing and novel cryptic pockets mapped across different simulation systems of the DENV E protein dimer. (A) Violin plot representing the distribution in SASA for the experimentally confirmed β-OG cryptic pocket during simulations, combining data from both chains and across all replicas. The pocket was reliably detected across all simulation systems except for sE_water_. Horizontal lines indicate the magnitude of apolar and polar surface areas measured for the pocket in the crystal structure^65^. (B) Violin plot representing the distribution in SASA of a novel cryptic site termed the “α pocket”, identified during simulations. The α pocket opened only during simulations in the presence of benzene. The pocket is predominantly hydrophobic, with a mean apolar SASA of 7.5 ± 1.6 nm^2^ and polar surface area of 5.5 ± 1.2 nm^2^ (dispersion is expressed as standard deviation). (C) A representative snapshot of the EM_bnz_-O simulation with the α and β-OG pockets in open states. The protein surface is coloured by electrostatic potential, ranging from -5 (red) to +5 (blue) kBT/e. The insets show the protein in a ribbon representation, coloured according to the functional region, including domain I (red), domain II (yellow), domain III (blue), and the stem and transmembrane helices (grey). The α pocket is located at the domain I – domain III interface, highlighted in lighter green. It is surrounded by the fusion loop and N^153^-glycan loop. The β-OG pocket occupies a space under the raised *kl* loop and is positioned at the domain I – domain II interface. The highlighted loops (fusion, N^153^-glycan, and *kl* loops) are shown as green ribbons in both insets.

All simulations in the presence of benzene successfully revealed the β-OG pocket, but EM_bnz_-P did so with the best agreement with values calculated for the crystal structure^65^ (mean apolar SASA determined to be 3.8 ± 1.5 nm^2^ and polar SASA 2.5 ± 1.0 nm^2^, compared to the crystallographic values of 3.3 and 2.7 nm^2^, respectively). The sE_bnz_ system, on the other hand, displayed a pocket accessibility in lesser agreement with experimental values (SASA_apolar_ = 4.0 ± 1.0 nm^2^; SASA_polar_ = 2.0 ± 0.6 nm^2^). Overexposure of the hydrophobic portion of the pocket suggests that the addition of benzene led to partial unfolding of the protein structure, expanding the cryptic pocket into the hydrophobic core of the protein which, in the native state, should never be exposed to the surrounding solvent.

#### 3.2.4 A novel pocket on the domain I – domain III interface

Novel cryptic pocket discovery was carried out by comparing pockets in systems containing benzene with control systems in a water-only environment. The biggest increase in pocket content was observed at the flexible domain I - domain III interface of the E protein, where a patch of unstructured surface loops is able to accommodate for benzene probe binding to the ectodomain interior. Strikingly, this is in close proximity to the hydrophobic fusion loop of the neighbouring chain, which must become exposed in order to anchor the viral particle to endosomal membranes and catalyze membrane fusion^38^. In this region, we consistently observed the formation of a cryptic pocket which we term the “α pocket”, which “snakes” through the domain I – domain III interface and reaches the fusion loop of a neighbouring chain (Fig. 6C). This α pocket was present in all simulations containing benzene, but was absent from the control runs (Fig. 6B). Measurement of its SASA shows that it is predominantly hydrophobic, consistent with its cryptic nature, which is also corroborated by the fact that benzene was able to transiently bind into the pocket groove. The residues surrounding the pocket involve residues at the N-terminus of the protein, the fusion loop, and the N^153^-glycan loop (Fig. S4). Interestingly, the interface of the two domains also contains a number of charged residues that are for the most part conserved across the representative flaviviral strains and which form localized patches of higher charge content within the pocket.

#### 3.2.5 Potential druggability of the α pocket

The α pocket exhibits a number of physicochemical characteristics that make it an interesting potential drug-binding hotspot: its elongated shape and a distinct pattern of charged and uncharged sites might contribute to drug-binding specificity; it is relatively large (2.20 ± 0.51 nm^3^) and therefore potentially able to establish a higher number of interactions with a drug-like molecule^66^; and the ionizable residues responsible for defining its charge distribution are conserved, suggesting that all dengue strains could simultaneously be targeted with the same α pocket-binding modulatory ligand.

Importantly, the portion of the protein where the α pocket is located undergoes a major conformational change triggered by the low-pH environment of the endosome^67^. This pocket is located at the hinge motion region between domain I and III and is important during the dimer-to-trimer transformation^38^. This process is crucial for the fusion of the viral and endosomal membranes and for subsequent release of the viral nucleocapsid into the cellular cytoplasm. A drug that is able to bind at the α pocket and stabilize both the domain I – domain III hinge region and the fusion loop would have the potential to act as an entry inhibitor, and to bind to the mature viral particle while it is still in the extracellular environment. This type of drug should remain in the bound state in both neutral and low-pH conditions, or else it would become ineffective during endocytosis. Notably, a drug-like molecule would also have an advantage over dengue-specific antibodies and vaccines because it would not trigger the antibody-dependent enhancement effects associated with the worst forms of dengue disease^41^.

Flavivirus particles are now known to undergo temperature-dependent expansion^37,68^, which may also be reversible depending on the surrounding salt conditions^69^, and their effects upon the stability of the α pocket remain to be explored. Since the pocket is formed by both E protein chains in the dimeric form, there is a possibility that the cryptic pocket could lose its defining shape in the expanded viral particle or during virion “breathing”. Similarly, the prM protein associated with the fusion loop in the immature virus (constituting around 30-40% of all secreted viral particles) might favour the closed conformation of a pocket, making it inaccessible to any pocket-specific drug molecule^70,71^. Also, in our simulated systems we do not take into consideration the impact of E protein glycans on the stability of the glycan loop. Likewise, potential interactions between the glycans and the surface pocket residues might play a role in shifting the conformational equilibrium of the pocket towards either closed or open state. These aspects therefore remain to be tested in future investigations for antiviral therapeutics development.

## 4 CONCLUSIONS

We have shown that our MD-based benzene mapping method can be successfully used for detection of cryptic pockets on membrane-bound proteins, as exemplified by proteins associated with the DENV envelope, if a repulsive term between benzene probes and specific lipid atoms is incorporated into the force field. Structural and dynamic properties of proteins embedded within a membrane are better matched with the conditions present in an enveloped viral particle when compared to the oversimplified ectodomain-only systems. MD simulations with benzene and lipid repulsion points on either P and O21/O31 atoms are most effective at revealing cryptic pockets, but EM_bnz_-O is more efficient at keeping the probes outside the bilayer. The method has proven successful in revealing the β-OG pocket that has been experimentally detected within the DENV E protein. We also uncovered a novel α pocket located on the domain I – domain III interface which extends to the fusion loop and has the potential to be a druggable site targeted by molecules acting as DENV entry inhibitors. The significance of the α pocket next needs to be assessed in the context of its structural consistency across a range of DENV serotypes and viral conformations occurring in the extracellular environment, as well in its ability to bind drug-like molecules capable of blocking the early stages of the DENV life cycle.

## 5 ACKNOWLEDGEMENTS

This work was supported by BII (A*STAR), the A*STAR Graduate Academy (A*GA), and the University of Manchester. PJB and JKM acknowledge NRF (NRF2017NRF-CRP001-027) for funding. The National Supercomputing Centre Singapore (NSCC) and the A*STAR Computational Resource Centre (A*CRC) provided computational resources. We thank Roland G. Huber for valuable discussions.

## REFERENCES

(1) Overington, J.; Hopkins, A. How Many Drug Targets Are There? Nat. Rev. Drug Discov. Drug Discov. 2006, 5 (December), 993–996. https://doi.org/10.1038/nrd2199.

(2) Abi Hussein, H.; Geneix, C.; Petitjean, M.; Borrel, A.; Flatters, D.; Camproux, A.-C. Global Vision of Druggability Issues: Applications and Perspectives. Drug Discov. Today 2017, 22 (2), 404–415. https://doi.org/10.1016/j.drudis.2016.11.021.

(3) Hajduk, P. J.; Huth, J. R.; Tse, C. Predicting Protein Druggability. Drug Discov. Today 2005, 10 (23–24), 1675–1682. https://doi.org/10.1016/S1359-6446(05)03624-X.

(4) Delano, W. L. Unraveling Hot Spots in Binding Interfaces: Progress and Challenges. Curr. Opin. Struct. Biol. 2002, 12, 14–20.

(5) Allen, K. N.; Bellamacina, C. R.; Ding, X.; Jeffery, C. J.; Mattos, C.; Petsko, G. A.; Ringe, D. An Experimental Approach to Mapping the Binding Surfaces of Crystalline Proteins †. J. Phys. Chem. 1996, 100 (7), 2605–2611. https://doi.org/10.1021/jp952516o.

(6) Ghanakota, P.; Carlson, H. A. Driving Structure-Based Drug Discovery through Cosolvent Molecular Dynamics. J. Med. Chem. 2016, 59, 10383–10399. https://doi.org/10.1021/acs.jmedchem.6b00399.

(7) Goodford, P. J. A Computational Procedure for Determining Energetically Favorable Binding Sites on Biologically Important Macromolecules. J. Med. Chem. 1985, 28, 849–857. https://doi.org/10.1021/jm00145a002.

(8) Miranker, A.; Karplus, M. Functionality Maps of Binding Sites: A Multiple Copy Simultaneous Search Method. Proteins Struct. Funct. Bioinforma. 1991, 11 (1), 29–34. https://doi.org/10.1002/prot.340110104.

(9) Brenke, R.; Kozakov, D.; Chuang, G. Y.; Beglov, D.; Hall, D.; Landon, M. R.; Mattos, C.; Vajda, S. Fragment-Based Identification of Druggable “hot Spots” of Proteins Using Fourier Domain Correlation Techniques. Bioinformatics 2009, 25 (5), 621–627. https://doi.org/10.1093/bioinformatics/btp036.

(10) Vigil, D.; Cherfils, J.; Rossman, K. L.; Der, C. J.Ras Superfamily GEFs and GAPs: Validated and Tractable Targets for Cancer Therapy? Nat. Rev. Cancer 2010, 10 (12), 842–857. https://doi.org/10.1038/nrc2960.

(11) Bowman, G. R.; Geissler, P. L.Equilibrium Fluctuations of a Single Folded Protein Reveal a Multitude of Potential Cryptic Allosteric Sites. Proc. Natl. Acad. Sci. 2012, 109 (29), 11681–11686. https://doi.org/10.1073/pnas.1209309109.

(12) Shaw, D. E.; Grossman, J. P.; Bank, J. A.; Batson, B.; Butts, J. A.; Chao, J. C.; Deneroff, M. M.; Dror, R. O.; Even, A.; Fenton, C. H.; et al. Anton 2: Raising the Bar for Performance and Programmability in a Special-Purpose Molecular Dynamics Supercomputer. Int. Conf. High Perform. Comput. Networking, Storage Anal. SC 2014, *2015*-*Janua* (January), 41–53. https://doi.org/10.1109/SC.2014.9.

(13) Lopes, P. E. M.; Guvench, O.; MacKerell, A. D.Current Status of Protein Force Fields for Molecular Dynamics Simulations. In *Molecular Modeling of Proteins*; Kukol, A., Ed.; Springer New York: New York, NY, 2015; pp 47–71. https://doi.org/10.1007/978-1-4939-1465-4_3.

(14) Durrant, J. D.; Keränen, H.; Wilson, B. A.; McCammon, J. A.Computational Identification of Uncharacterized Cruzain Binding Sites. PLoS Negl. Trop. Dis. 2010, 4 (5), 1–11. https://doi.org/10.1371/journal.pntd.0000676.

(15) Johnson, D. K.; Karanicolas, J.Druggable Protein Interaction Sites Are More Predisposed to Surface Pocket Formation than the Rest of the Protein Surface. PLoS Comput. Biol. 2013, 9 (3). https://doi.org/10.1371/journal.pcbi.1002951.

(16) Oleinikovas, V.; Saladino, G.; Cossins, B. P.; Gervasio, F. L.Understanding Cryptic Pocket Formation in Protein Targets by Enhanced Sampling Simulations. J. Am. Chem. Soc. 2016, 138, 14257–14263. https://doi.org/10.1021/jacs.6b05425.

(17) Seco, J.; Luque, F. J.; Barril, X.Binding Site Detection and Druggability Index from First Principles. J. Med. Chem. 2009, 52 (8), 2363–2371.

(18) Tan, Y. S.; Reeks, J.; Brown, C. J.; Thean, D.; Ferrer Gago, F. J.; Yuen, T. Y.; Goh, E. T. L.; Lee, X. E. C.; Jennings, C. E.; Joseph, T. L.; et al. Benzene Probes in Molecular Dynamics Simulations Reveal Novel Binding Sites for Ligand Design. J. Phys. Chem. Lett. 2016, 7 (17), 3452–3457. https://doi.org/10.1021/acs.jpclett.6b01525.

(19) Bakan, A.; Nevins, N.; Lakdawala, A. S.; Bahar, I.Druggability Assessment of Allosteric Proteins by Dynamics Simulations in the Presence of Probe Molecules. J. Chem. Theory Comput. 2012, 8 (7), 2435–2447. https://doi.org/10.1021/ct300117j.

(20) Kimura, S. R.; Hu, H. P.; Ruvinsky, A. M.; Sherman, W.; Favia, A. D.Deciphering Cryptic Binding Sites on Proteins by Mixed-Solvent Molecular Dynamics. J. Chem. Inf. Model. 2017, 57 (6), 1388–1401. https://doi.org/10.1021/acs.jcim.6b00623.

(21) Guvench, O.; MacKerell, A. D.Computational Fragment-Based Binding Site Identification by Ligand Competitive Saturation. PLoS Comput. Biol. 2009, 5 (7), e1000435. https://doi.org/10.1371/journal.pcbi.1000435.

(22) Lexa, K. W.; Carlson, H. A.Full Protein Flexibility Is Essential for Proper Hot-Spot Mapping. J. Am. Chem. Soc. 2011, 133 (2), 200–202. https://doi.org/10.1021/ja1079332.

(23) Sayyed-Ahmad, A.; Gorfe, A. A.Mixed-Probe Simulation and Probe-Derived Surface Topography Map Analysis for Ligand Binding Site Identification. J. Chem. Theory Comput. 2017, 13 (4), 1851–1861. https://doi.org/10.1021/acs.jctc.7b00130.

(24) Schames, J. R.; Henchman, R. H.; Siegel, J. S.; Sotriffer, C. A.; Ni, H.; McCammon, J. A.Discovery of a Novel Binding Trench in HIV Integrase. J. Med. Chem. 2004, 47, 1879–1881. https://doi.org/10.1021/jm0341913.

(25) De Young, L. R.; Dill, K. A.Solute Partitioning into Lipid Bilayer Membranes. Biochemistry 1988, 27 (14), 5281–5289. https://doi.org/10.1021/bi00414a050.

(26) Yildirim, M. A.; Goh, K.; Cusick, M. E.; Barabási, A.; Vidal, M.Drug—Target Network. Nat. Biotechnol. 2007, 25 (10), 1119–1126. https://doi.org/10.1038/nbt1338.

(27) Prakash, P.; Sayyed-Ahmad, A.; Gorfe, A. A.PMD-Membrane: A Method for Ligand Binding Site Identification in Membrane-Bound Proteins. PLoS Comput. Biol. 2015, 11 (10), 1–20. https://doi.org/10.1371/journal.pcbi.1004469.

(28) Landon, M. R.; Amaro, R. E.; Baron, R.; Ngan, C. H.; Ozonoff, D.; Andrew McCammon, J.; Vajda, S.Novel Druggable Hot Spots in Avian Influenza Neuraminidase H5N1 Revealed by Computational Solvent Mapping of a Reduced and Representative Receptor Ensemble. Chem. Biol. Drug Des. 2008, 71 (2), 106–116. https://doi.org/10.1111/j.1747-0285.2007.00614.x.

(29) Abdelnabi, R.; Geraets, J. A.; Ma, Y.; Mirabelli, C.; Flatt, J. W.; Domanska, A.; Delang, L.; Jockhmans, D.; Kumar, T. A.; Jayaprakash, V.; et al. A Novel Druggable Interprotomer Pocket in the Capsid of Rhino- and Enteroviruses. PLOS Biol. 2019, 71, 1–17.

(30) Lok, S.-M. The Interplay of Dengue Virus Morphological Diversity and Human Antibodies. Trends Microbiol. 2016, 24 (4), 284–293. https://doi.org/10.1016/j.tim.2015.12.004.

(31) Mukhopadhyay, S.; Kuhn, R. J.; Rossmann, M. G.A Structural Perspective of the Flavivirus Life Cycle. Nature Reviews Microbiology. 2005, pp 13–22. https://doi.org/10.1038/nrmicro1067.

(32) Gould, E. A.; Solomon, T.Pathogenic Flaviviruses. Lancet 2008, 371, 500–509. https://doi.org/10.1590/S0034-72802009000200010.

(33) Liu-Helmersson, J.; Quam, M.; Wilder-Smith, A.; Stenlund, H.; Ebi, K.; Massad, E.; Rocklöv, J.Climate Change and Aedes Vectors: 21st Century Projections for Dengue Transmission in Europe. EBioMedicine 2016, 7, 267–277. https://doi.org/10.1016/j.ebiom.2016.03.046.

(34) Bhatt, S.; Gething, P. W.; Brady, O. J.; Messina, J. P.; Farlow, A. W.; Moyes, C. L.; Drake, J. M.; Brownstein, J. S.; Hoen, A. G.; Sankoh, O.; et al. The Global Distribution and Burden of Dengue. Nature 2013, 496 (7446), 504–507. https://doi.org/10.1038/nature12060.

(35) Zhang, X.; Ge, P.; Yu, X.; Brannan, J. M.; Bi, G.; Zhang, Q.; Schein, S.; Hong Zhou, Z.Cryo-EM Structure of the Mature Dengue Virus at 3.5-Å Resolution. Nat. Struct. Mol. Biol. 2013, 20 (1), 105–110. https://doi.org/10.1038/nsmb.2463.

(36) Fibriansah, G.; Ibarra, K. D.; Ng, T. S.; Smith, S. A.; Tan, J. L.; Lim, X. N.; Ooi, J. S. G.; Kostyuchenko, V. A.; Wang, J.; De Silva, A. M.; et al. Cryo-EM Structure of an Antibody That Neutralizes Dengue Virus Type 2 by Locking E Protein Dimers. Science (80-.). 2015, 349 (6243), 88–91. https://doi.org/10.1126/science.aaa8651.

(37) Zhang, X.; Sheng, J.; Plevka, P.; Kuhn, R. J.; Diamond, M. S.; Rossmann, M. G.Dengue Structure Differs at the Temperatures of Its Human and Mosquito Hosts. Proc. Natl. Acad. Sci. 2013, 110 (17), 6795–6799. https://doi.org/10.1073/pnas.1304300110.

(38) Modis, Y.; Ogata, S.; Clements, D.; Harrison, S. C.Structure of the Dengue Virus Envelope Protein after Membrane Fusion. Nature 2004, 427 (6972), 313–319. https://doi.org/10.1038/nature02165.

(39) Yu, I.-M.; Zhang, W.; Holdaway, H. A.; Li, L.; Kostyuchenko, V. A.; Chipman, P. R.; Kuhn, R. J.; Rossmann, M. G.; Chen, J.Structure of the Immature Dengue Virus at Low PH Primes Proteolytic Maturation. Science (80-.). 2008, 319 (5871), 1834–1837. https://doi.org/10.1126/science.1153264.

(40) Li, L.; Lok, S. M.; Yu, I. M.; Zhang, Y.; Kuhn, R. J.; Chen, J.; Rossmann, M. G.The Flavivirus Precursor Membrane-Envelope Protein Complex: Structure and Maturation. Science (80-.). 2008, 319 (5871), 1830–1834. https://doi.org/10.1126/science.1153263.

(41) Katzelnick, L. C.; Gresh, L.; Halloran, M. E.; Mercado, J. C.; Kuan, G.; Gordon, A.; Balmaseda, A.; Harris, E.Antibody-Dependent Enhancement of Severe Dengue Disease in Humans. Science (80-.). 2017, 358, 929–932.

(42) Wirawan, M.; Fibriansah, G.; Marzinek, J. K.; Crowe, J. E.; Bond, P. J.; Wirawan, M.; Fibriansah, G.; Lim, X. X.; Ng, T.; Sim, A. Y. L.; et al. Mechanism of Enhanced Immature Dengue Virus Attachment to Endosomal Membrane Induced by PrM Antibody Article Mechanism of Enhanced Immature Dengue Virus Attachment to Endosomal Membrane Induced by PrM Antibody. Struct. Des. 2019, 27, 1–15. https://doi.org/10.1016/j.str.2018.10.009.

(43) Huang, J.; Mackerell, A. D.CHARMM36 All-Atom Additive Protein Force Field: Validation Based on Comparison to NMR Data. J. Comput. Chem. 2013, 34 (25), 2135–2145. https://doi.org/10.1002/jcc.23354.

(44) Hess, B., Bekker, H., Berendsen, H.J.C. & Fraaije, J. G. E. M. LINCS: A Linear Constraint Solver for Molecular Simulations. Journal of Computational Chemistry. J. Comput. Chem. 1997, 18 (18), 1463–1472. https://doi.org/10.1002/(SICI)1096-987X(199709)18:12<1463::AID-JCC4>3.0.CO;2-H.

(45) Soni, A.K. Uncovering Cryptic Pockets in Biologically Relevant Proteins: An Improved Computational Methodology. Master’s Thesis, Indian Institute of Science Education and Research Pune, Maharashtra, India. 2017.

(46) Zhang, Q.; Hunke, C.; Yau, Y. H.; Seow, V.; Lee, S.; Tanner, L. B.; Guan, X. L.; Wenk, M. R.; Fibriansah, G.; Chew, P. L.; et al. The Stem Region of Premembrane Protein Plays an Important Role in the Virus Surface Protein Rearrangement during Dengue Maturation. J. Biol. Chem. 2012, 287 (48), 40525–40534. https://doi.org/10.1074/jbc.M112.384446.

(47) Sunhwan, J.; Taehoon, K.; Vidyashankara, I. G.; Wonpil, I.CHARMM-GUI: A Web-Based Graphical User Interface for CHARMM. J. Comput. Chem. 2008, 29, 1859–1865. https://doi.org/10.1002/jcc.

(48) Abraham, M. J.; Murtola, T.; Schulz, R.; Páll, S.; Smith, J. C.; Hess, B.; Lindah, E.Gromacs: High Performance Molecular Simulations through Multi-Level Parallelism from Laptops to Supercomputers. SoftwareX 2015, 1–2, 19–25. https://doi.org/10.1016/j.softx.2015.06.001.

(49) Jorgensen, W. L.; Chandrasekhar, J.; Madura, J. D.; Impey, R. W.; Klein, M. L.Comparison of Simple Potential Functions for Simulating Liquid Water. J. Chem. Phys. 1983, 79 (2), 926–935. https://doi.org/10.1063/1.445869.

(50) MacKerell, A. D.; Bashford, D.; Bellott, M.; Dunbrack, R. L.; Evanseck, J. D.; Field, M. J.; Fischer, S.; Gao, J.; Guo, H.; Ha, S.; et al. All-Atom Empirical Potential for Molecular Modeling and Dynamics Studies of Proteins. J. Phys. Chem. B 1998, 102 (18), 3586–3616. https://doi.org/10.1021/jp973084f.

(51) Lomize, M. A.; Pogozheva, I. D.; Joo, H.; Mosberg, H. I.; Lomize, A. L.OPM Database and PPM Web Server: Resources for Positioning of Proteins in Membranes. Nucleic Acids Res. 2012, 40 (D1), 370–376. https://doi.org/10.1093/nar/gkr703.

(52) Wu, E. L.; Cheng, X.; Jo, S.; Rui, H.; Song, K. C.; Dávila-Contreras, E. M.; Qi, Y.; Lee, J.; Monje-galvan, V.; Venable, R. M.; et al. CHARMM-GUI Membrane Builder Toward Realistic Biological Membrane Simulations. J. Comput. Chem. 2015, 35 (27), 1997–2004. https://doi.org/10.1002/jcc.23702.CHARMM-GUI.

(53) Bussi, G.; Donadio, D.; Parrinello, M.Canonical Sampling through Velocity Rescaling Canonical Sampling through Velocity Rescaling. J. Chem. Phys. 2007, 126, 014101. https://doi.org/10.1063/1.2408420.

(54) Parrinello, M.; Rahman, A.Polymorphic Transitions in Single Crystals: A New Molecular Dynamics Method. J. Appl. Phys. 1981, 52 (12), 7182–7190. https://doi.org/10.1063/1.328693.

(55) Essmann, U.; Perera, L.; Berkowitz, M. L.; Darden, T.; Lee, H.; Pedersen, L. G.A Smooth Particle Mesh Ewald Method. J. Chem. Phys. 1995, 103 (19), 8577–8593. https://doi.org/10.1063/1.470117.

(56) Eisenhaber, F.; Lijnzaad, P.; Argos, P.; Sander, C.; Scharf, M.The Double Cubic Lattice Method: Efficient Approaches to Numerical Integration of Surface Area and Volume and to Dot Surface Contouring of Molecular Assemblies. J. Comput. Chem. 1995, 16 (3), 273–284. https://doi.org/10.1002/jcc.540160303.

(57) Kabsch, W.; Sander, C.Dictionary of Protein Secondary Structure: Pattern Recognition of Hydrogen-bonded and Geometrical Features. Biopolymers 1983, 22 (12), 2577–2637. https://doi.org/10.1002/bip.360221211.

(58) Buchoux, S.FATSLiM: A Fast and Robust Software to Analyze MD Simulations of Membranes. Bioinformatics 2017, 33 (1), 133–134. https://doi.org/10.1093/bioinformatics/btw563.

(59) Schmidtke, P.; Bidon-Chanal, A.; Luque, F. J.; Barril, X.MDpocket : Open Source Cavity Detection and Characterization on Molecular Dynamics Trajectories. Bioinformatics 2011, 1–10.

(60) Jurrus, E.; Engel, D.; Star, K.; Monson, K.; Brandi, J.; Felberg, L. E.; Brookes, D. H.; Wilson, L.; Chen, J.; Liles, K.; et al. Improvements to the APBS Biomolecular Solvation Software Suite. Protein Sci. 2018, 27 (1), 112–128. https://doi.org/10.1002/pro.3280.

(61) Porter, J. R.; Moeder, K. E.; Sibbald, C. A.; Zimmerman, M. I.; Hart, K. M.; Greenberg, M. J.; Bowman, G. R.Cooperative Changes in Solvent Exposure Identify Cryptic Pockets, Switches, and Allosteric Coupling. Biophys. J. 2019, 116 (5), 818–830. https://doi.org/10.1016/j.bpj.2018.11.3144.

(62) Lim, X.-X.; Chandramohan, A.; Lim, X. Y. E.; Bag, N.; Sharma, K. K.; Wirawan, M.; Wohland, T.; Lok, S.-M.; Anand, G. S.Conformational Changes in Intact Dengue Virus Reveal Serotype-Specific Expansion. Nat. Commun. 2017, 8, e14339. https://doi.org/10.1038/ncomms14339.

(63) Cheng, B.; Wu, S.; Liu, S.; Rodriguez-Aliaga, P.; Yu, J.; Cui, S.Protein Denaturation at a Single-Molecule Level: The Effect of Nonpolar Environments and Its Implications on the Unfolding Mechanism by Proteases. Nanoscale 2015, 7 (7), 2970–2977. https://doi.org/10.1039/c4nr07140a.

(64) Hosseinzadeh, R.; Moosavi-Movahedi, A. A.Human Hemoglobin Structural and Functional Alterations and Heme Degradation upon Interaction with Benzene: A Spectroscopic Study. Spectrochim. Acta - Part A Mol. Biomol. Spectrosc. 2016, 157, 41–49. https://doi.org/10.1016/j.saa.2015.12.014.

(65) Modis, Y.; Ogata, S.; Clements, D.; Harrison, S. C.A Ligand-Binding Pocket in the Dengue Virus Envelope Glycoprotein. Proc. Natl. Acad. Sci. 2003, 100 (12), 6986–6991. https://doi.org/10.1073/pnas.0832193100.

(66) Hajduk, P. J.; Huth, J. R.; Fesik, S. W.Druggability Indices for Protein Targets Derived from NMr-Based Screening Data. J. Med. Chem. 2005, 48 (7), 2518–2525. https://doi.org/10.1021/jm049131r.

(67) Stiasny, K.; Allison, S. L.; Marchler-Bauer, A.; Kunz, C.; Heinz, F. X.Structural Requirements for Low-PH-Induced Rearrangements in the Envelope Glycoprotein of Tick-Borne Encephalitis Virus. J. Virol. 1996, 70 (11), 8142–8147.

(68) Fibriansah, G.; Ng, T.-S.; Kostyuchenko, V. A.; Lee, J.; Lee, S.; Wang, J.; Lok, S.-M. Structural Changes in Dengue Virus When Exposed to a Temperature of 37 C. J. Virol. 2013, 87 (13), 7585–7592. https://doi.org/10.1128/JVI.00757-13.

(69) Sharma, K. K.; Lim, X. X.; Tantirimudalige, S. N.; Gupta, A.; Marzinek, J. K.; Holdbrook, D.; Lim, X. Y. E.; Bond, P. J.; Anand, G. S.; Wohland, T.Infectivity of Dengue Virus Serotypes 1 and 2 Is Correlated with E-Protein Intrinsic Dynamics but Not to Envelope Conformations. Structure 2019, 27 (4), 618-630.e4. https://doi.org/10.1016/j.str.2018.12.006.

(70) van der Schaar, H. M.; Rust, M. J.; Waarts, B.-L.; van der Ende-Metselaar, H.; Kuhn, R. J.; Wilschut, J.; Zhuang, X.; Smit, J. M.Characterization of the Early Events in Dengue Virus Cell Entry by Biochemical Assays and Single-Virus Tracking. J. Virol. 2007, 81 (21), 12019–12028. https://doi.org/10.1128/jvi.00300-07.

(71) Zybert, I. A.; van der Ende-Metselaar, H.; Wilschut, J.; Smit, J. M.Functional Importance of Dengue Virus Maturation: Infectious Properties of Immature Virions. J. Gen. Virol. 2008, 89 (12), 3047–3051. https://doi.org/10.1099/vir.0.2008/002535-0.

