## Supporting Information for "A benzene-mapping approach for uncovering cryptic pockets in membrane-bound proteins"

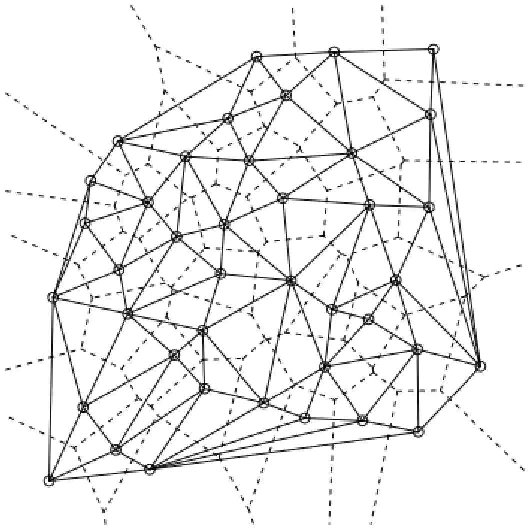

**Figure S1. Voronoi vertices representing each lipid by its phosphorous atom coordinate.** Circles represent P atom positions in a single leaflet based on its *xy*-coordinates (because of the planar nature of the membrane, the *z*-coordinate is disregarded). Solid lines connecting the neighbouring lipids are a measure of lateral distance. For each lipid, only the longest distance is considered. A Voronoi diagram does not take into consideration periodic boundary conditions, resulting in an incorrect neighbour mapping in case of edge vertices. The edge vertices are therefore excluded from the calculation, except for their role as neighbours to non-edge lipids.

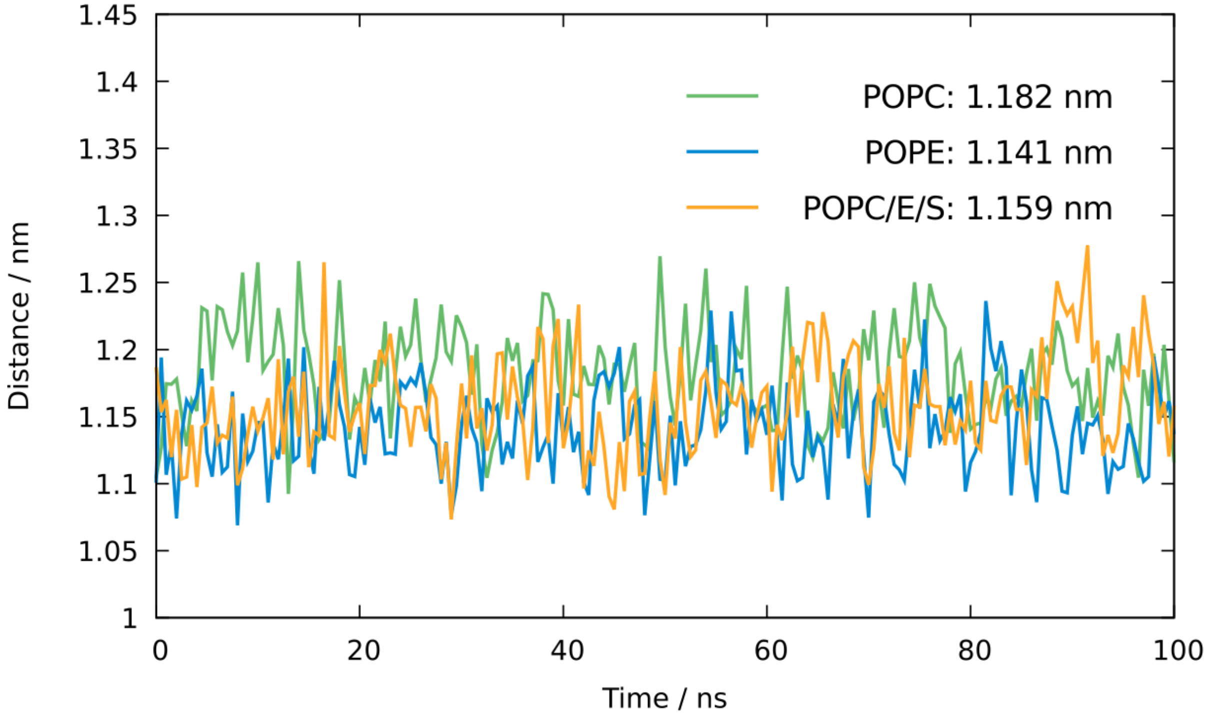


**Figure S2. Time progression of the maximum lateral gap between a lipid and its neighbours.** The mean distance value is specified for each membrane type. The values are averaged across all non-edge lipids in both leaflets. The membranes shown are simulated only in water, without benzene present in the system.

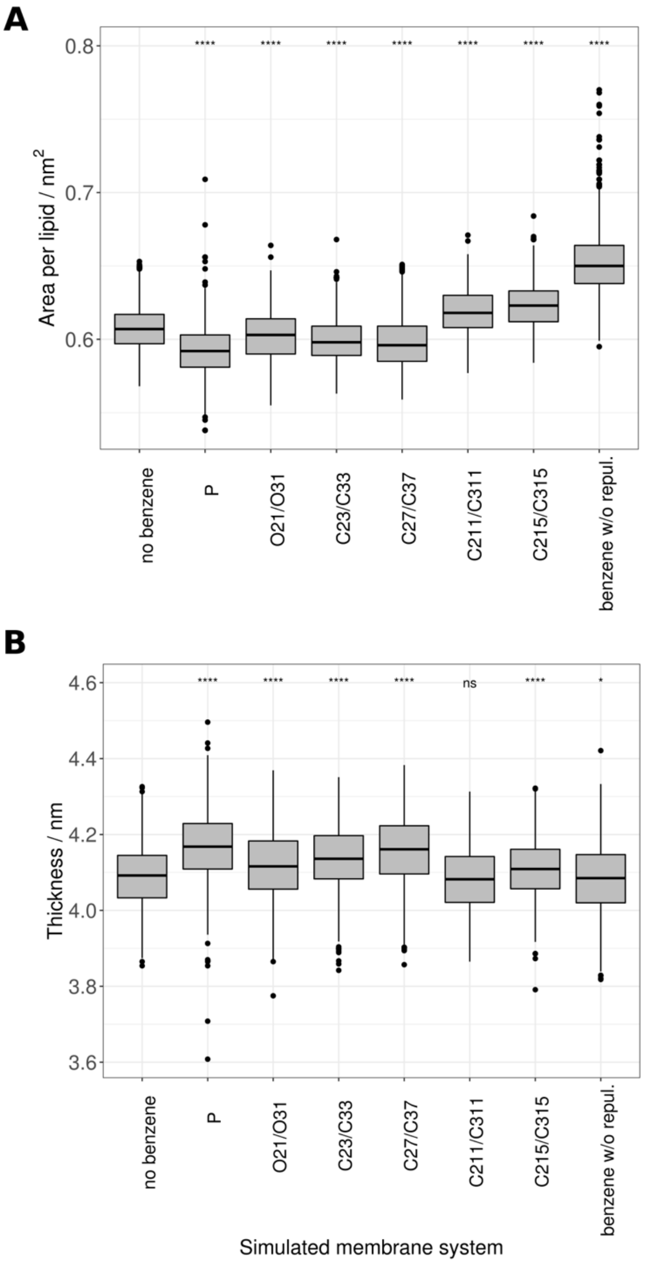


**Figure S3. Membrane properties depending on the placement of the lipid atom repulsion point.** All simulations refer to the heterogeneous DENV membrane containing a mixture of three phospholipid types. (A) Area per lipid measurement is affected by the presence of benzene in the systems, with the biggest difference observed in a system containing benzene molecules but without repulsive forces imposed. (B) The effect of benzene on membrane thickness is less obvious, C211/C311 being the only system not showing a significant difference in the means. Statistical analysis was performed using Student's *t*-test with all systems being compared against the reference control group (*no benzene). ns:* $p > 0.5$; * : $p \leq 0.05$; **** : $p \leq 0.0001$.

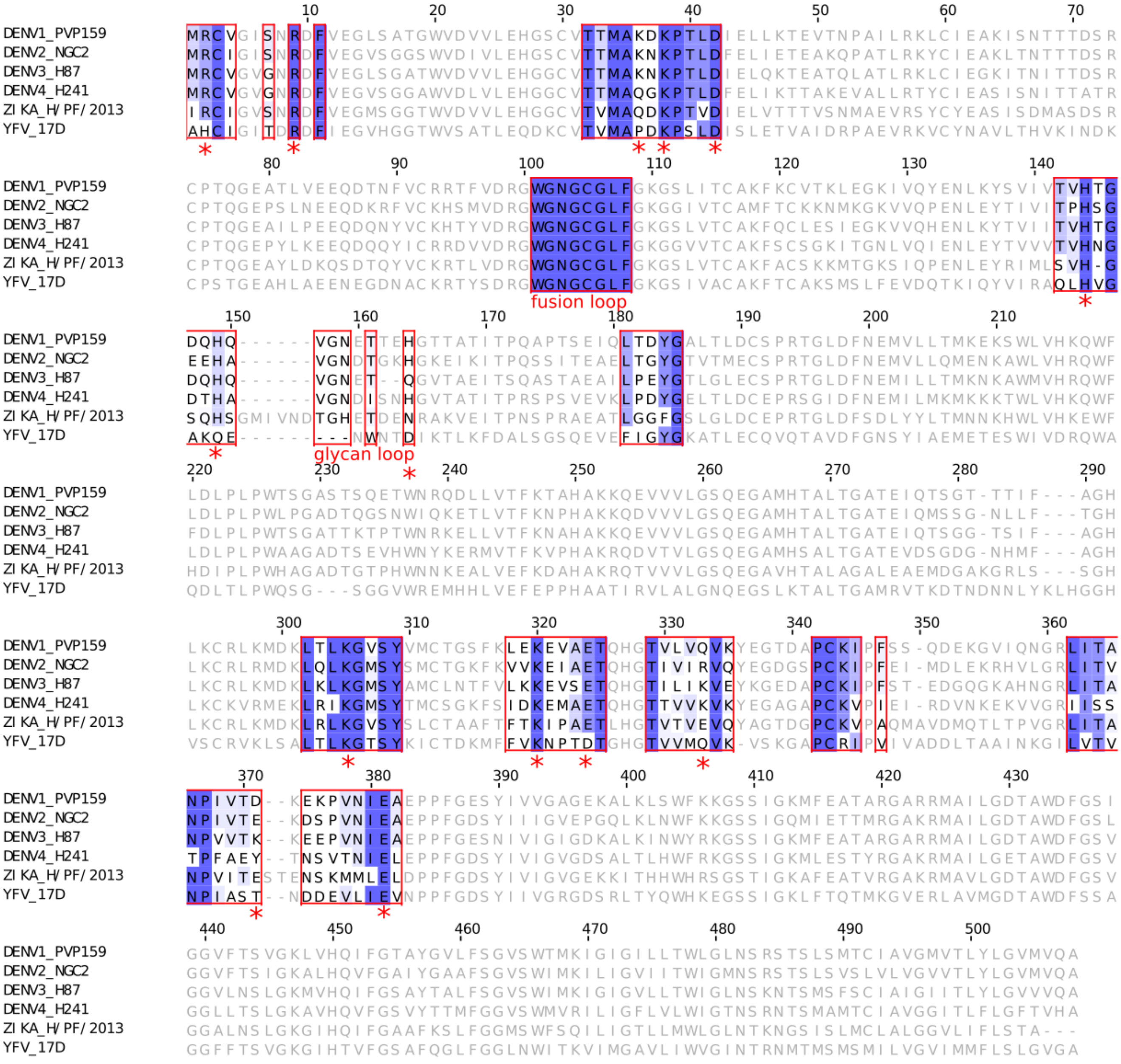


**Figure S4. Multiple alignment of the E-protein showing the residues in the vicinity of the α pocket (highlighted in red boxes).** Residues are coloured according to their conservation within the representative flaviviral strains. The pocket contains a number of residues belonging to domain I and domain III, as well as the domain II fusion loop (labelled). Ionisable residues, if present in the DENV-2 sequence, are marked with red stars and are, for the most part, conserved across the viral family.
